# Genomic data reveal a north-south split and introgression history of blood fluke (*Schistosoma haematobium*) populations from across Africa

**DOI:** 10.1101/2024.08.06.606828

**Authors:** Roy N Platt, Egie E. Enabulele, Ehizogie Adeyemi, Marian O Agbugui, Oluwaremilekun G Ajakaye, Ebube C Amaechi, Chika E Ejikeugwu, Christopher Igbeneghu, Victor S Njom, Precious Dlamini, Grace A. Arya, Robbie Diaz, Muriel Rabone, Fiona Allan, Bonnie Webster, Aidan Emery, David Rollinson, Timothy J.C. Anderson

## Abstract

The human parasitic fluke, *Schistosoma haematobium* hybridizes with the livestock parasite *S. bovis* in the laboratory, but the frequency of hybridization in nature is unclear. We analyzed 34.6 million single nucleotide variants in 162 samples from 18 African countries, revealing a sharp genetic discontinuity between northern and southern *S. haematobium*. We found no evidence for recent hybridization. Instead the data reveal admixture events that occurred 257-879 generations ago in northern *S. haematobium* populations. Fifteen introgressed *S. bovis* genes are approaching fixation in northern *S. haematobium* with four genes potentially driving adaptation. We identified 19 regions that were resistant to introgression; these were enriched on the sex chromosomes. These results (i) suggest strong barriers to gene flow between these species, (ii) indicate that hybridization may be less common than currently envisaged, but (iii) reveal profound genomic consequences of rare interspecific hybridization between schistosomes of medical and veterinary importance.

## Introduction

Hybridization and the transfer of alleles via introgression is an important source of genetic variation between species^1^. This process allows for allelic variants, which have already been preselected in a donor species, to be introduced into the genome of a recipient species in a single generation. By comparison, it may take multiple generations for random mutation and selection to deliver comparable levels of genetic variation in the absence of introgression^2^. As a result, introgressive hybridization, can contribute to the evolution of new genetic traits in hybridizing species^3^. Hybridization between human and animal pathogens can lead to the emergence of parasites with novel traits such as increased pathogenicity^4^, expanded host range^5^, altered transmission dynamics^6^ and drug resistance^7^. Understanding the frequency and impact of such hybridization events is critical for devising effective disease intervention strategies.

Members of the blood fluke genus *Schistosoma* parasitize a range of mammal species and cause substantial morbidity and economic loss^8^. Parasites in this genus have a complex life cycle: first-stage larvae (miracidia) infect an aquatic snail intermediate host where they develop into sporocysts. Clonally generated, motile second-stage larvae (cercariae) emerge from the intermediate host, and actively locate and penetrate the skin of the definitive mammalian host. In the mammal host, mature parasites form male-female pairs in the blood stream and reproduce sexually. Eggs are excreted through the feces or urine, depending on the parasite species, which restarts the life cycle.

One pair of species, *S. haematobium*, a human parasite, and *S. bovis*, an ungulate parasite common in domestic livestock, are genetically divergent (3-5% and 18% divergent in the nuclear and mitochondrial genomes respectively), but can hybridize and produce viable offspring when given the opportunity^9^. Given the close proximity between humans and their livestock and the regular use of the same water bodies, the potential for hybridization between these species is a particular concern and a significant effort has been mounted to identify, monitor and map *S. haematobium* and *S. bovis* hybrids^10^. Multiple studies have reported mitochondrial and/or ribosomal DNA from *S. bovis* in *S. haematobium* populations. For examples see ^9,11–13^. The high frequency of individuals with discordant mitochondrial and nuclear markers has been used to argue that hybridization is common and that the zoonotic threat of *S. bovis* should be considered in human schistosomiasis control programs^14^.

However, several, more recent multi-marker and genomic studies using single nucleotide variants (SNVs), microsatellite markers and whole genome sequence assemblies have suggested that hybridization between *S. haematobium* and *S. bovis* may not occur as frequently as previously postulated. Exome^15^, whole genome^16^, and microsatellite data^17^, and others^18–21^ failed to identify evidence of contemporary hybridization in field-collected parasites. Instead, these studies indicate that *S. bovis* and *S. haematobium* are genetically distinct and do not hybridize frequently but evidence for historical hybridization is clearly evident within genomes of *S. haematobium*. As a result, some *S. bovis* genes have introgressed into the *S. haematobium* population and reached high frequency; evidence of a potential, adaptive introgression event^15,16,18,20^.

In this study, we build upon previous work and try to address knowledge gaps by analyzing a comprehensive dataset of 34.6 million genome-wide SNVs from *S. bovis* (n=21) and *S. haematobium* (n=141) samples collected from 18 countries across the African continent. Many of these samples presented discordant mitochondrial and ribosomal DNA profiles and were categorized as *S. haematobium*-*bovis* hybrids. This expanded dataset, and recent availability of a high-quality *S. haematobium* genome assembly^22^, allows for a more detailed examination of the genetic relationships between these two species and the potential consequences of hybridization on their evolution and epidemiology. Our results: (i) reveal a strong discontinuity between northern and southern *S. haematobium* populations; (ii) define similar genomic introgression profiles in *S. haematobium* sampled from locations 3,002 Km apart; (iii) fine-map introgressed genome regions and identify putative genes driving adaptive introgression; (iv) identify two distinct lineages of *S. bovis*-like mitochondrial DNA in northern *S. haematobium*, consistent with historical introgression and (v) identify “introgression deserts” on the ZW chromosomes consistent with the sex chromosomes maintaining species integrity. These results enhance our understanding of *Schistosoma spp.* epidemiology, with important implications for control efforts.

## Results

### DNA Sequencing and Genotyping

We examined 219 *Schistosoma* samples from 24 countries (Figure 1A). Just over 80% (n=176) of the samples were collected as part of this study with the remaining 43 samples made available through open access resources. Median genome coverage per sample was 29.7x. After filtering, the final dataset contained 35,817,757 total SNVs, 7,206,957 common (minor allele frequency; MAF>0.05%) SNVs, and 446,162 common unlinked SNVs genotyped from 162 samples (141 *S. haematobium* and 21 *S. bovis*; Figure 1B). Filtered samples (n=51) were removed due to low genome coverage (mean = 9.7x) compared to passing samples (mean=30.8x). NCBI Short Read Archive (SRA) accessions and sample metadata are available in Supplemental Table 1.

**Figure 1.**
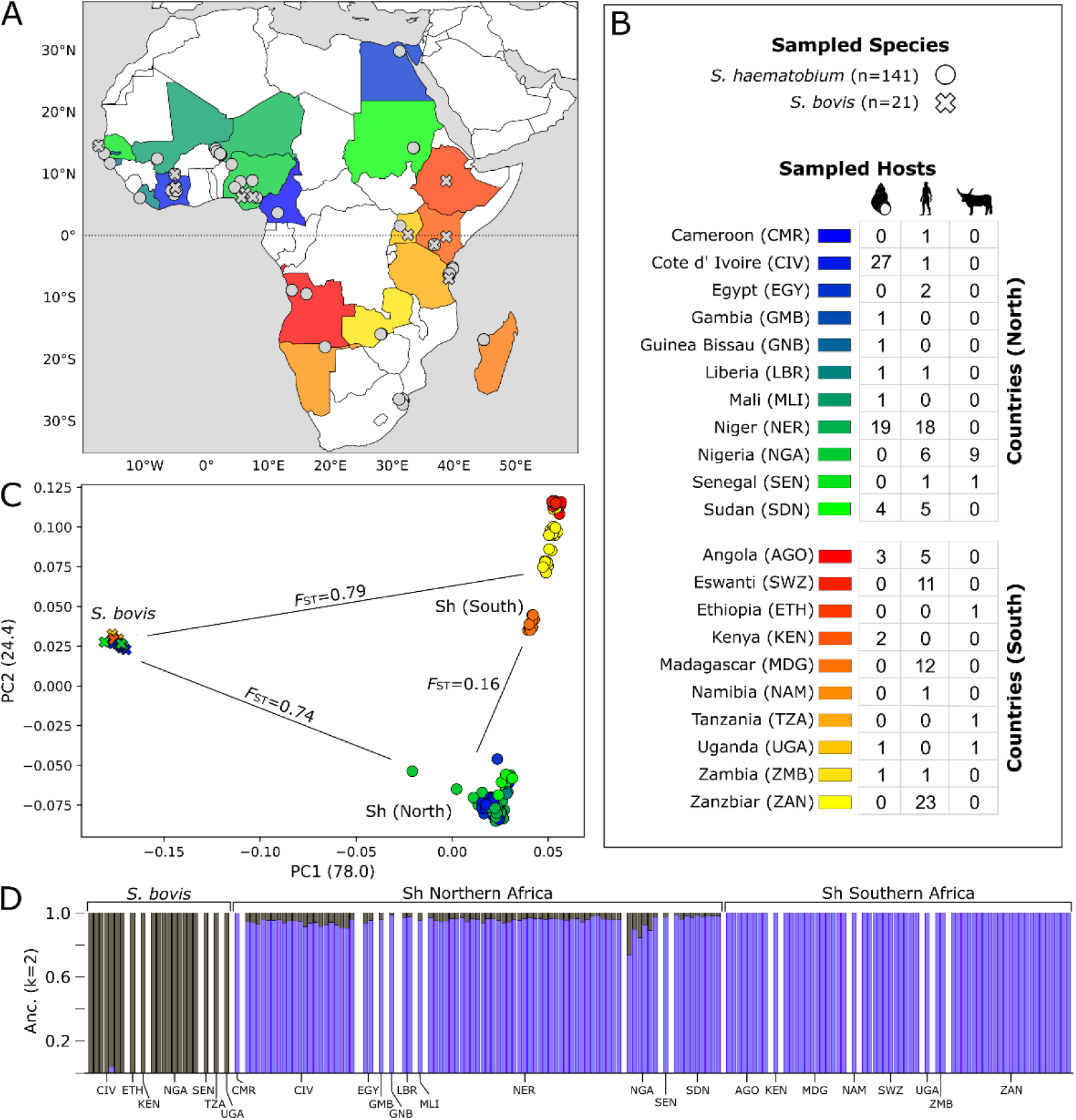
Sampling localities, sample summary and the population structure of *Schistosoma haematobium* and *S. bovis*. (A) Collection locations for samples used in this study. Where exact coordinates for samples were not readily available we used the country capital as the collection locality. Two populations of *S. haematobium* were identified; northern and southern. The southern population in red-yellow and the northern population in blue to green. (B) A description of species, hosts, and countries sampled in this study. (C) A principal component analysis of 355,715 unlinked, common (MAF>0.05), autosomal variants. The three clusters correspond to *S. bovis,* and the northern and southern *S. haematobium* populations. Weighted, Weir-Cockerham *F*ST values between these populations are shown. (D) A supervised admixture analysis (k=2) was used to assign ancestry to each sample. This analysis shows almost all of the northern *S. haematobium* samples contained low levels of *S. bovis*. Country Codes are as follows: “AGO”: Angola, “CMR”: Cameroon, “CIV”: Cote d’ Ivoire, “EGY”: Egypt, “SWZ”: Eswanti, “ETH”: Ethiopia, “GMB”: Gambia, “GNB”: Guinea Bissau, “KEN”: Kenya, “LBR”: Liberia, “MDG”: Madagascar, “MLI”: Mali, “NAM”: Namibia, “NER”: Niger, “NGA”: Nigeria, “SEN”: Senegal, “SDN”: Sudan, “TZA”: Tanzania, “UGA”: Uganda, “ZMB”: Zambia, “ZAN”: Zanzibar.

### Population structure and ancestry

We examined relationships among samples with a PCA of 355,715 unlinked, common, autosomal SNVs (Figure 1C). Each of our samples fell into one of 3 *K*-means clusters along PC1 and PC2. The three clusters corresponded with (a) *S. haematobium* individuals from northern Africa, (b) southern Africa, and (c) all *S. bovis* samples. The northern population includes samples collected in Cameroon, Cote d’ Ivoire, Egypt Gambia, Guinea Bissau, Liberia, Mali, Niger, Nigeria, Senegal and Sudan. The southern population includes samples collected from Angola, Eswatini, Kenya, Madagascar, Namibia, Tanzania, Uganda, Zambia, and Zanzibar. In general, the equator approximately delineates the northern and southern populations. The division of *S. haematobium* into northern and southern populations was consistent among analyses with one exception. Madagascar was an intermediate population in Admixture analysis (k=3) (Supplemental Figure 2). In the PCA, samples from Madagascar area assigned to the southern cluster, but they form a distinct subgroup that is intermediate between the remaining southern and northern samples.

No samples were placed intermediate between the *S. haematobium* and *S. bovis* clusters which would indicate F1 *S. haematobium*-*bovis* hybrids among these samples. The weighted, Weir and Cockerham *F*ST^23,24^ between the *S. bovis* and *S. haematobium* samples was high (*F*ST ≥ 0.74-0.79). We observed strong subdivision between northern and SE *S. haematobium* populations (*F*ST = 0.16; Figure 1C) with multiple *F*ST peaks (Figure 2D). There were 275,657 SNVs showing fixed differences (*F*ST = 1) between *S. bovis* and *S. haematobium* (Supplemental Table 2). Mean sequence divergence (*d_XY_*) between *S. haematobium* and *S. bovis* was 0.015 compared to 0.002 between the northern and southern *S. haematobium* populations (Supplemental Figure 1).

**Figure 2.**
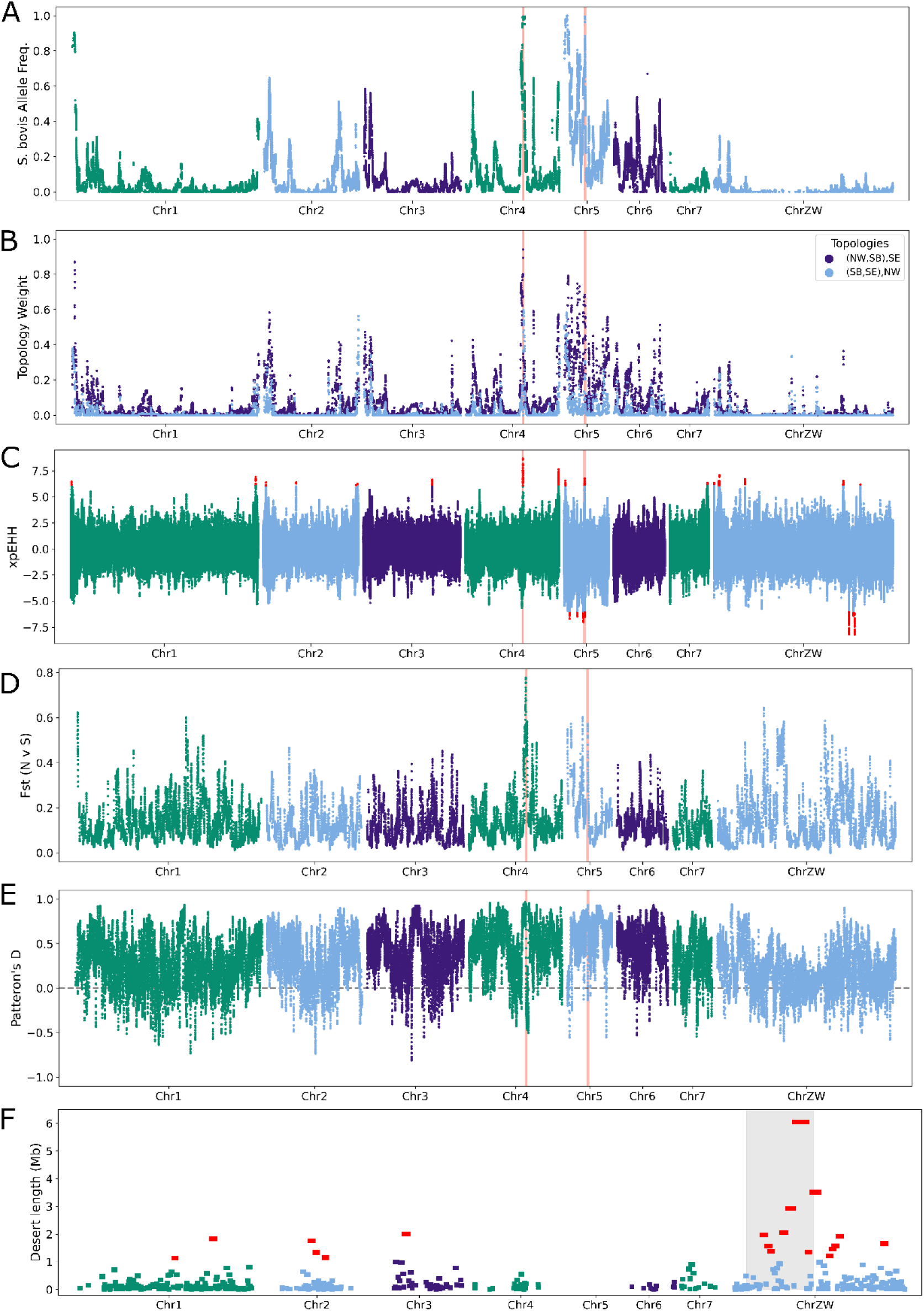
Local measurements of differentiation, introgression and selection across the genome. (A) The frequency of *S. bovis* ancestry across the genome in the northern *S. haematobium* population was estimated using RFmix. While the percentage of *S. bovis* alleles in the population is low overall, the *S. bovis* alleles are at or near fixation at loci on Chr4 and Chr5. (B) Gene tree topology weightings across the genome depicting the possible relationships between the northern and south *S. haematobium* populations and *S. bovis* using TWISST. Each locus across the genome is shown as stacked bar plots. While both tools use different methods to depict the relationships between these taxa they recover similar results. (C) Differential selection between *S. haematobium* populations was measured across the genome with extended haplotype homozygosity (xpEHH). Positive values indicate positive selection in the northern population and negative values indicate positive selection in the southern population. Significant xpEHH values (*p*<0.05) after multiple test correction are highlighted in red. (D) The weighted Weir-Cockerham fixation index (*F*ST) between northern and southern Africa, S. haematobium populations was measured across the genome in 10Kb windows. These results indicated multiple, highly differentiated regions between the two populations. (E) Patterson’s *D* statistic was measured to determine if high *F*ST regions were the result of introgressed *S. bovis* alleles present in northern populations using a test tree of (((southern *S*. *haematobium*, northern *S. haematobium*), *S. bovis*), *S. margrebowiei*). *D* measured across the genome was significantly positive indicating the presence of gene flow between *S. bovis* and north African *S. haematobium* populations. (F) Multiple regions of the northern African *S. haematobium* genome lacked introgressed *S. bovis* alleles. Introgression desserts that are longer than expected by chance are shown (Z-scoreLength > 3) in red. The grey box represents the Z-specific region of the sex chromosome. Results for FST and Patterson’s D are shown after Gaussian smoothing (sigma=3). Pink vertical lines indicate putative regions of adaptive introgression.

We used Admixture v1.3.0^25^ to quantify ancestry among the samples (Figure 1D, Supplemental Figure 2). We found that when *k* =2, *S. bovis* and south African *S. haematobium* individuals were exclusively assigned with different ancestry components. By contrast *S. haematobium* samples collected from northern Africa were a composite of the two population components including 0.5-26.2% (median = 4.2%) of the *S. bovis* population component (Supplemental Figure 3).

### Reference biases

We tested the data for read-mapping reference biases, which could occur if non-*haematobium* species map poorly to the Egyptian-strain *S. haematobium* reference assembly (GCF_000699445.3), which groups with the northern population in our analyses. Mapping rates were 77.7%, 82.8% and 76.3% for *S. bovis*, northern *S. haematobium* and southern *S. haematobium* populations (Figure 1A). A *t*-Test failed to identify differential mapping rates between *S. haematobium* and *S. bovis* (*p* = 0.397), suggesting that reference bias does not significantly contribute to the results observed. The modest mapping rate results from the complexity of the genome which contains high levels of repetitive elements^26^.

### Phylogenetics

The species tree generated using SVDquartets and nuclear SNVs revealed a well-resolved topology. We examined 2,500,000 random quartets, representing 8.82% of all possible distinct quartets. Of these, 18.5% (n = 463,571) were incompatible with the final tree (Figure 3). *S. haematobium* and *S. bovis* were resolved into two clades and *S. haematobium* individuals fell into clades reflecting geographic relationships.

**Figure 3.**
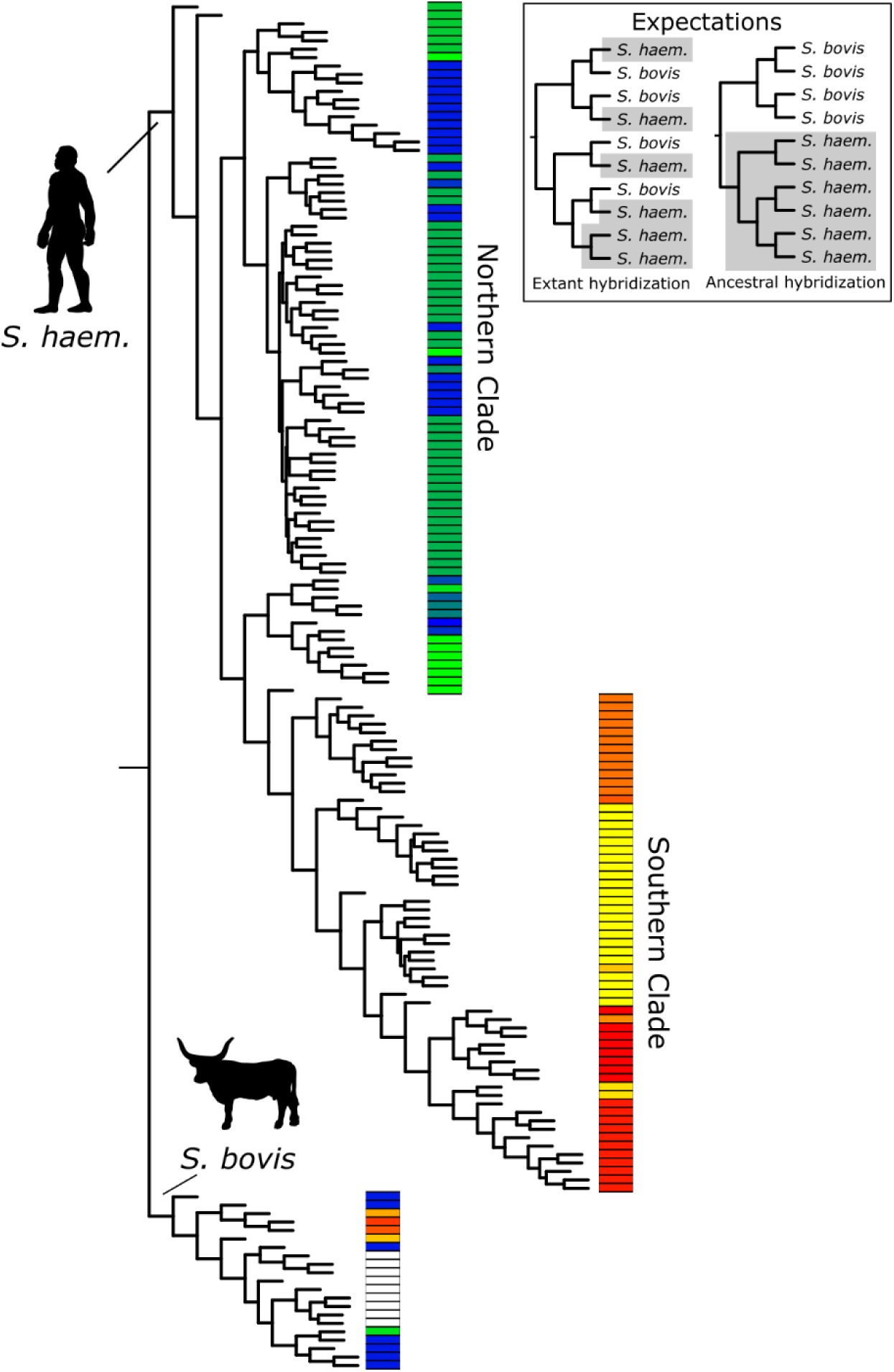
Species tree of *S. haematobium* and *S. bovis* populations. SVDquartets species tree generated from autosomal SNVs. All nodes were supported by >95% of bootstrap replicates. Phylogenetic relationships between the species can be used to differentiate extant vs ancestral hybridization (inset). The tree shows that both *S. haematobium* and *S. bovis* are monophyletic. Biogeographic partitioning within the tree indicates that *S. haematobium* originated in northern Africa and expanded into southern Africa.

On average, we were able to assemble 15,558.4 bp of the mitochondrial genome for each sample, 4,757 of which were phylogenetically informative sites. The mitochondrial phylogeny (Figure 4) reveals two major mitochondrial haplotypes, one containing *S. haematobium* individuals from across Africa and another clade containing all of the *S. bovis* and 38 north African *S. haematobium*. The presence of *S. haematobium* samples within a larger *S. bovis* clade is consistent with *S. bovis* mitochondrial introgression into *S. haematobium* that has been frequently reported in field samples, for example ^27^. Within the *S. bovis* clade, all *S. haematobium* samples with the introgressed *S. bovis* mitochondria fell into two monophyletic groups, clades “A” and “B”. mtDNA haplotypes from these two clades were from samples widely distributed in northern Africa. For example, the same clade “A” haplotype was found in samples from Egypt, Niger and Cote d’ Ivoire (>3,300 km apart). The clade “B” haplotype was found in Niger, Nigeria and Cote d’ Ivoire, a linear distance of 1,171 km. Bootstrap support for each of these major clades was strong (100%). Phylogenetic trees in Newick format are available in the supplementary materials.

**Figure 4.**
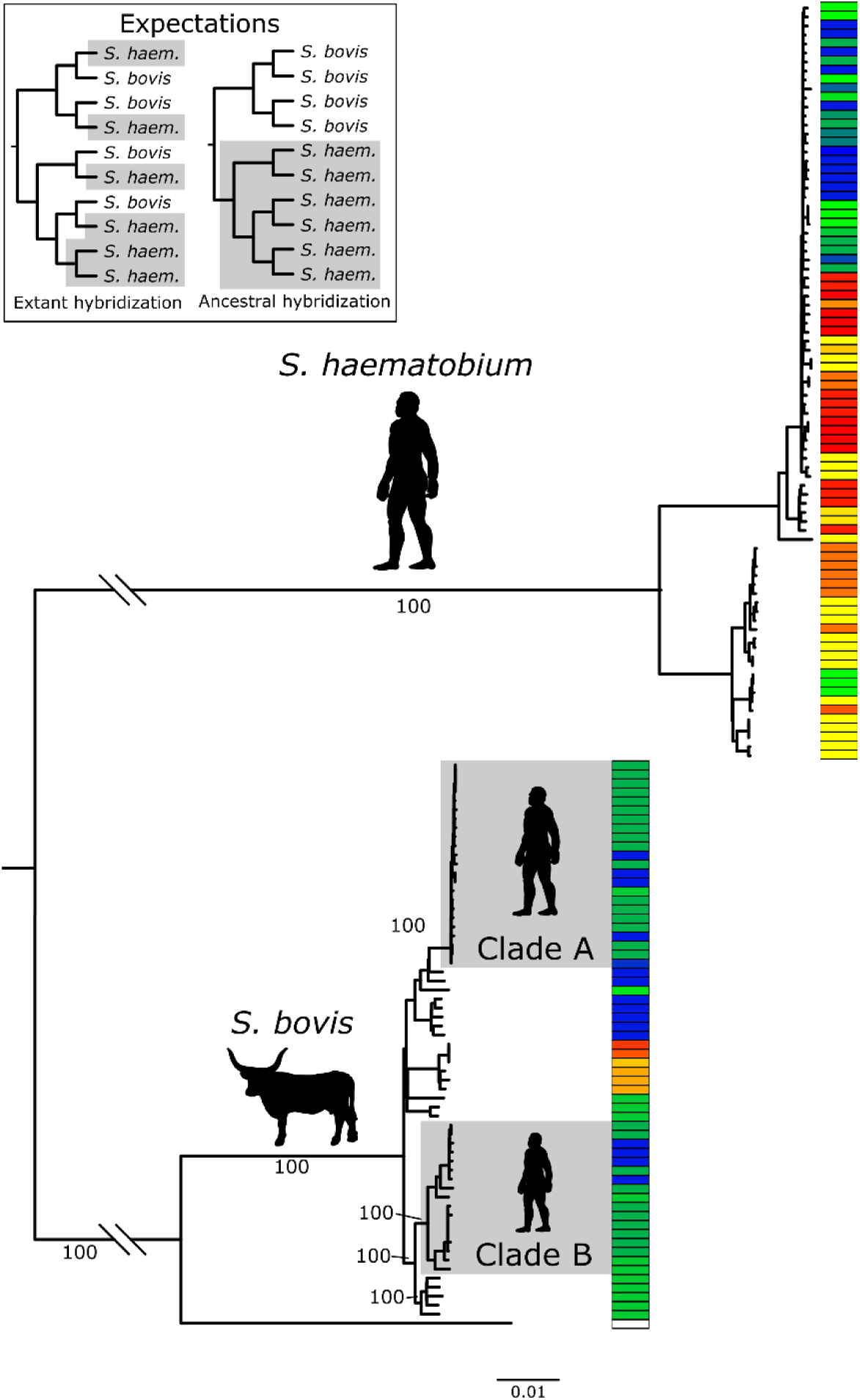
Mitochondrial tree of *S. haematobium* and *S. bovis*. A gene tree was recovered from mitochondrial genome assemblies from each sample. Bootstrap support at select nodes is shown. Phylogenetic relationships between the species can be used to differentiate extant vs ancestral hybridization (inset). Two well supported clades of *S. haematobium* contain an introgressed *S. bovis* mitotype, designated as “A” and “B”. Both the “A” and “B” clades contain samples from north Africa. All south African samples are found within a single clade of the remaining *S. haematobium* samples

### Hybridization and Introgression

We used four methods to identify signatures of hybridization and introgression between *S. haematobium* and *S. bovis*. These methods include *f_3_*, *D-*statistic, local ancestry assignment (RFmix^28^) and phylogenetic discordance (TWISST^29^). First, a negative *f_3_* (N: S, Sb; *f_3_ =*-0.128, SE= 0.8*e*^-3^, z-score=-156.4) indicates that north African *S. haematobium* population include *S. bovis* ancestry. Next, we used the *D*-statistic to test for introgression between *S. haematobium* and *S. bovis* while accounting for lineage sorting (Figure 2E). We averaged *D* in 10Kb blocks. *D* was significantly positive (*D*=0.46, σ_M_=0.007, n=30,278) indicating biased introgression between north African *S. haematobium* and *S. bovis*.

We examined the landscape of introgression across the genome using local ancestry with RFmix. We used 38 southern African *S. haematobium* lacking *S. bovis* introgression and 13 *S. bovis* samples to serve as reference panels for “pure” *S. haematobium* and *S. bovis*. RFmix results showed that ancestry across the genome was not uniformly distributed in the north African population (Figure 2A). Within the north African population, *S. bovis* ancestry blocks ranged in frequency from 0-100% at any particular locus. Each north African *S. haematobium* sample contained 4.1-22.0% *S. bovis* ancestry (median 7.0%). By comparison the median *S. bovis* ancestry was 0.02% and 100% in the southern *S. haematobium* and *S. bovis* control samples, respectively.

We used TWISST v67b9a66^29^ as an independent method for identifying local introgression. TWISST measures shifting gene tree frequencies across the genome. Trees were generated from 37,200 non-overlapping, 10kb, sliding windows. On average (mean) each window contained 657.3 SNVs. We examined the three possible topologies between northern *S. haematobium*, southern *S. haematobium* and *S. bovis* using *S. margrebowiei* (GCA_944470205.2^30^) as an outgroup (Figure 2B). The expected species tree, with a monophyletic clade of *S. haematobium*, sister to *S. bovis* was the most common with a mean weight of 0.876 across the genome. The discordant topology uniting northern *S. haematobium* and *S. bovis* was the second most abundant topology (weight = 0.085) compared to the topology with southern *S. haematobium* and *S. bovis* (weight = 0.039).

We examined the genome for regions that are devoid of introgressed *S. bovis* alleles in the northern *S. haematobium* population. There were 918 genomic deserts lacking *S. bovis* alleles with a median size of 35.8Kb (Figure 2F). With log transformation and robust Z-scores we identified 19 genomic deserts that were significant outliers in terms of length ranging from 1.13-6 Mb (median 1.67 Mb). Thirteen of the 19 deserts were on the ZW scaffold and accounted for 32% (28.6 Mb) of its total length.

### Introgression profiles in different countries

We examined the pattern of introgression in individual countries of north Africa (Supplemental Figure 4) as determined by RFMix. The overall patterns of introgression across the genome were consistent between north African populations. Pairwise comparisons of introgressed allele frequencies between countries were positively correlated (*r* = 0.59-0.8; Supplemental Figure 5) despite distances spanning up to 3,000 km.

### Impact of introgression on nucleotide diversity and genetic differentiation of *S. haematobium*

We masked introgressed alleles within individual genomes, and recalculated π, *F*ST and a PCA (Figure 5). Prior to masking, mean nucleotide diversity (π), was 2.3-fold greater in the northern (π = 2.991 x 10^-3^) vs southern (π = 1.278 x 10^-3^) *S. haematobium*, and π was 3.3-fold greater in *S. bovis* (π = 8.329 x 10^-3^) than the entirety of *S. haematobium* (π = 2.523 x 10^-3^). After masking, northern African *S. haematobium* nucleotide diversity was reduced to nearly identical levels seen in the south population: π_NW_ = 2.991 x 10^-3^ to π_NW_ = 1.07 x 10^-3^. By comparison, removing introgressed alleles had minimal impact on *F*ST (*F*ST = 0.154) between northern and southern *S. haematobium.* Additionally, the structure of the PCA was retained, demonstrating that introgression makes a minimal contribution to genome-wide differentiation between northern and southern populations.

**Figure 5.**
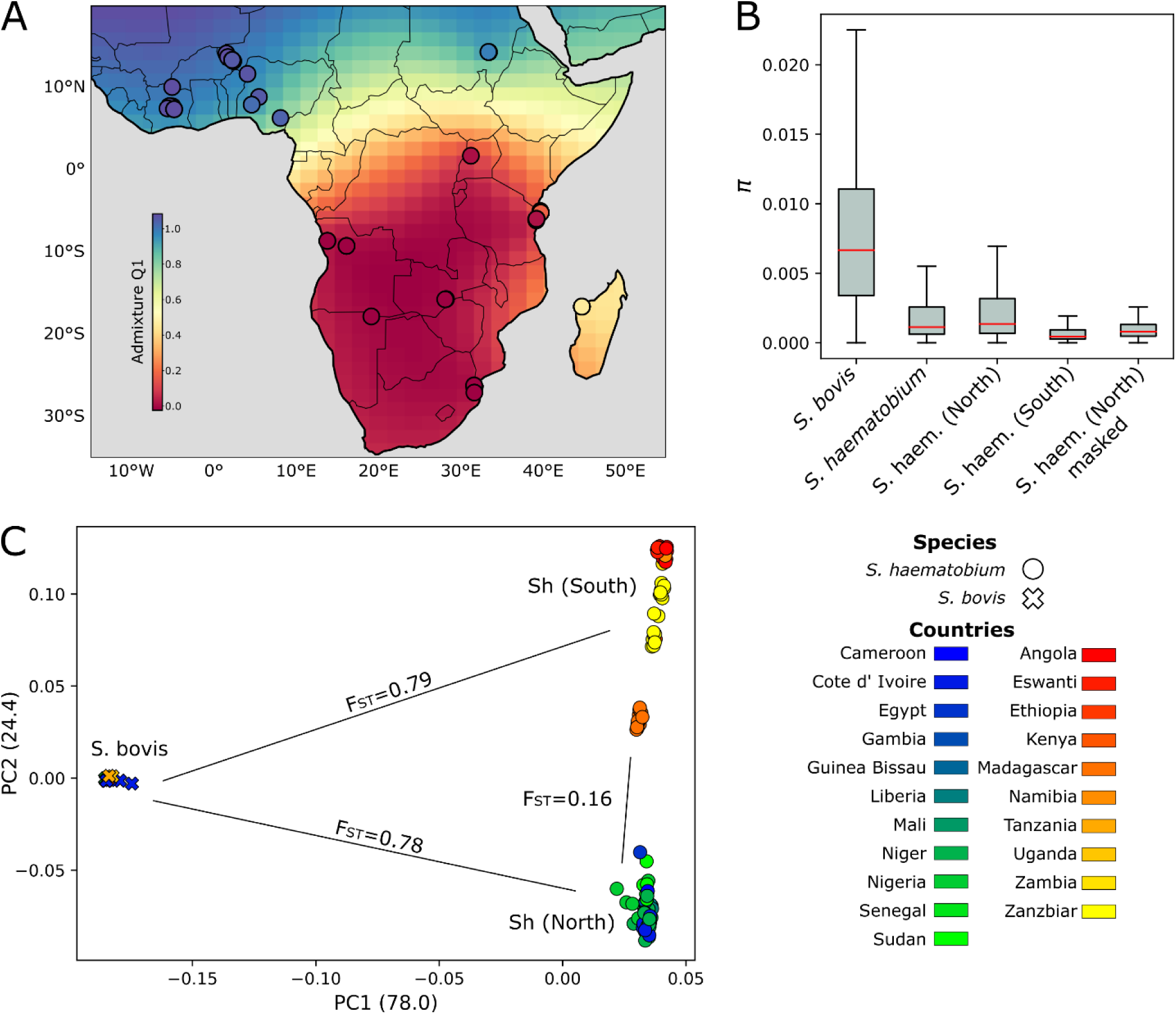
Biogeography of *S. haematobium* is not determined by introgressed *S. bovis* alleles. *S. haematobium* samples were split into two populations by PCA, Admixture and phylogenetic analyses. (A) We used Kriging interpolation to examine the distribution of these populations across Africa using the population component that differentiates the *S. haematobium* populations from one another (B) Nucleotide diversity (π) was calculated in 10kb sliding windows after masking introgressed *S. bovis* alleles present in the northern *S. haematobium population*. π is higher in the northern African *S. haematobium* compared to the southern population. When introgressed *S. bovis* alleles are masked, π is similar for both the southern and northern populations. (C) After masking introgressed *S. bovis* alleles the PCA is similar to Figure 1C. The similarity between the two PCAs show that the genetic differentiation between the northern and southern *S. haematobium* populations is not driven by introgressed *S. bovis* alleles.

### Dating Introgression

We used the size of introgressed haplotype blocks to estimate the number of generations since hybridization for each *S. haematobium* sample in north Africa (Supplemental Figure 6). This gave estimated hybridization dates of ∼257-879 generations ago (Median – 426 generations, 95% confidence intervals = 281.6-764 generations). *S. haematobium* generation time varies from 3-4^31^ months in lab populations, but is estimated to be 6-12 months in wild populations^32^. These generation times imply that admixture between *S. haematobium* and *S. bovis* occurred ∼106 years ago assuming four generations per year (high transmission) or 426 years assuming one generation per year (low and / or seasonal transmission). Dating estimates varied between countries: median estimates are lowest in Egypt (286.9) and highest in Nigeria (565) despite their relatively close proximity. A one-way ANOVA indicated significant differences in the number of generations since hybridization between countries (p-value = 1.3*e*^-10^; Supplemental Figure 6).

### Selection and adaptive introgression

We examined *S. haematobium* and *S. bovis* populations for signatures of selection using normalized, xpEHH (Figure 2C). We found 996 statistically significant xpEHH values after multiple test correction. We combined values within 1Mb to identify 15 genome regions with signatures of positive selection in the northern population and five in the southern population (Supplemental Table 3). The median normalized xpEHH in each of these regions was >|6| and the windows ranged in size from 3 bp to 709,928 bp (mean 139,942 bp).

We combined selection and introgression analyses to identify genome regions showing evidence for adaptive introgression of *S. bovis* alleles into the northern *S. haematobium* population. These regions contained outlier values for selection (xpEHH), elevated Patterson’s D (D ≥ 0) indicative of introgression, high levels of S. *bovis* ancestry (>95%) and significant differentiation from southern *S. haematobium* (*F*ST ≥ 95^TH^ percentile). Two genome regions met these criteria; chromosome four (NC_067199.1:28,476,500-28,813,500) and chromosome five (NC_067200.1:9,773,000-10,447,000). These genome regions span 1.01 Mb and 15 genes; eight on chr four and seven on chr five (Table 1). Of the 74,955 SNVs in these regions, 989 are nonsense or missense mutations. We found 37 missense SNVs where the *S. bovis* allele is at or near fixation in the northern population (Supplemental Table 4). All of these variants are on chr four and fall within four genes; leishmanolysin-like peptidase, a Rho GTPase-activating protein 35, Jumonji domain-containing protein six (JMJD6_4), and Jumonji domain-containing protein six (JMJD6_3).

**Table 1.**
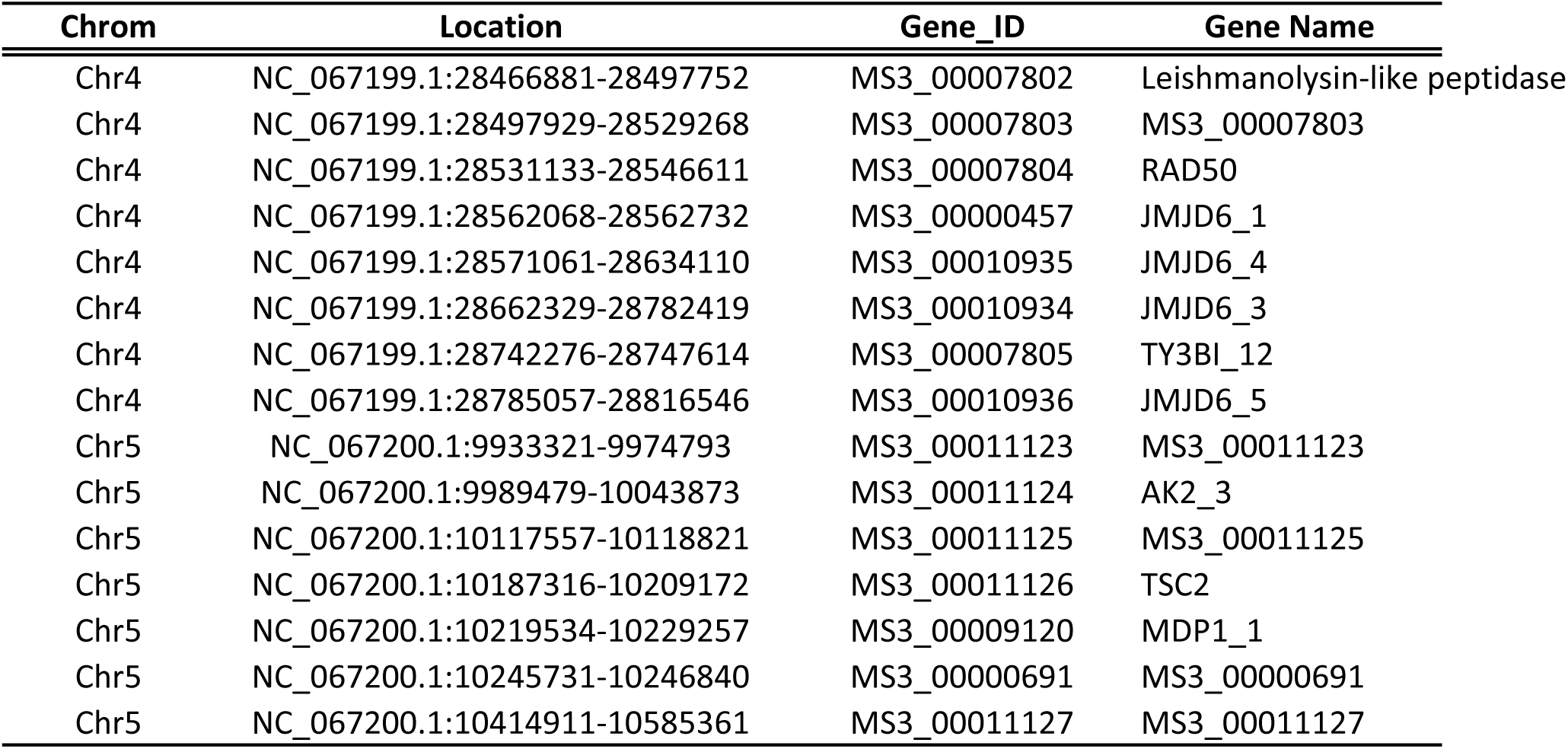
Genes containing outlier loci.

| **Chrom** | **Location** | **Gene_ID** | **Gene Name** |
| --- | --- | --- | --- |
| Chr4 | NC_067199.1:28466881-28497752 | MS3_00007802 | Leishmanolysin-like peptidase |
| Chr4 | NC_067199.1:28497929-28529268 | MS3_00007803 | MS3_00007803 |
| Chr4 | NC_067199.1:28531133-28546611 | MS3_00007804 | RAD50 |
| Chr4 | NC_067199.1:28562068-28562732 | MS3_00000457 | JMJD6_1 |
| Chr4 | NC_067199.1:28571061-28634110 | MS3_00010935 | JMJD6_4 |
| Chr4 | NC_067199.1:28662329-28782419 | MS3_00010934 | JMJD6_3 |
| Chr4 | NC_067199.1:28742276-28747614 | MS3_00007805 | TY3BI_12 |
| Chr4 | NC_067199.1:28785057-28816546 | MS3_00010936 | JMJD6_5 |
| Chr5 | NC_067200.1:9933321-9974793 | MS3_00011123 | MS3_00011123 |
| Chr5 | NC_067200.1:9989479-10043873 | MS3_00011124 | AK2_3 |
| Chr5 | NC_067200.1:10117557-10118821 | MS3_00011125 | MS3_00011125 |
| Chr5 | NC_067200.1:10187316-10209172 | MS3_00011126 | TSC2 |
| Chr5 | NC_067200.1:10219534-10229257 | MS3_00009120 | MDP1_1 |
| Chr5 | NC_067200.1:10245731-10246840 | MS3_00000691 | MS3_00000691 |
| Chr5 | NC_067200.1:10414911-10585361 | MS3_00011127 | MS3_00011127 |

## Discussion

Our analysis of >38 million SNVs provides compelling evidence that *S. haematobium* and *S. bovis* are genetically well differentiated. This conclusion is supported by multiple lines of evidence: high *F*ST values (*F*ST ≥ 0.74-0.79; Figure 1C; Figure 2D), distinction by PCA (Figure 1C), strong differentiation by ancestry components in Admixture analyses (Figure 1D) and well supported monophyletic clades in the nuclear species tree (Figure 3). The agreement between these approaches suggests that strong barriers to gene flow exist between these two species.

Our analysis revealed that northern African *S. haematobium* are genetically differentiated (*F*ST = 0.16) from the southern population. The boundary between these populations appears to extend from Cameroon, Gabon, the Central African Republic, South Sudan, and Ethiopia (Figure 5A) and approximately follows the equator. When introgressed *S. bovis* alleles were removed from genomic data, *F*ST between these populations is only marginally affected (*F*ST = 0.154). Hence, introgressed *S. bovis* alleles have a minimal impact on genetic differences between the northern and southern populations.

Our results suggest barriers to gene flow exist between northern and southern *S. haematobium*. The southern *S. haematobium* clade is nested within a larger clade of northern African *S. haematobium* (Figure 3). This indicates that *S. haematobium* originated in one of the northern African countries and is consistent with previous work that identified the Arabian Peninsula/Asia as a potential ancestral source population^33^. It is possible that the two populations are defined by the distribution of their intermediate hosts. Regional differences in parasite compatibility with their intermediate snail hosts can occur within limited geographical areas^34^. *S. haematobium* is primarily transmitted by members of the *Bulinus truncatus/tropicus* complex from North Africa and the Middle East are primarily transmitted by the *Bulinus truncatus*/*tropicus* species complex and parasites from the Afro-tropical region are transmitted by snails of the *Bulinus globosus* and member of the *africanus* species group^35^, although exceptions to this rule exist^36^. *S. bovis* by comparison is transmitted by many of the same snail species including members of the B. *truncatus*/*tropicus* and *africanus* species groups, and *B. forskalii*^21^. Other, less frequently studied species may influence these dynamics as well^37,38^. If a barrier exists that is related to the intermediate snail hosts, it has important implications for our understanding of the ecological and epidemiological factors that shape the distribution and dynamics of these two parasite populations. Further investigation at the population boundaries may provide new insights into biological differences and incompatibilities between northern and southern *S. haematobium* populations. We note that the existence of northern and southern populations of *S. haematobium* based on the use of snail intermediate hosts was suggested in the last century^39^.

Four aspects of our results support a historical introgression hypothesis. First, each of the north African *S. haematobium* samples contain low levels of *S. bovis* ancestry with the exception of the sole Cameroonian sample. Percentages of *S. bovis* ancestry per individual are similar across multiple analyses: introgressed haplotype blocks from RFMix account for 4.1-22% of individual genomes in the northern *S. haematobium* population, while the population component associated with *S. bovis* in Admixture ranges from 5-26.2% at *K*=2.

Second, the landscape of introgressed alleles across the genome is consistent across north African samples (Supplemental Figure 4) and positively correlated (Supplemental Figure 5) despite being separated by ≤3,000 Km. For this profile to be conserved across such a broad distance suggests (A) it occurred in an ancestor of the north African *S. haematobium* or (B) introgressed alleles provided a selective advantage that spread throughout the north African population. Our data support the later with the nuclear phylogeny (Figure 3). The southern population is a monophyletic clade that lacks introgressed *S. bovis* alleles, although the limitations of a bifurcating tree should be considered under scenarios of introgression. We also observe that some introgressed alleles have reached high frequency in the north African population and show signs of selection (Figure 2). Finally, our data suggest that there is a barrier to gene flow/migration between northern and southern *S. haematobium* populations, restricting dispersal of introgressed alleles to the southern population.

Third, mitochondrial DNA provides insights into a minimal number of hybridization events. 58% of northern *S. haematobium* samples contain introgressed S*. bovis* mitochondria (Figure 4). If the introgressed *S. bovis* mitochondria were the result of contemporary hybridization, we would expect sister relationships between *S. bovis* and *S. haematobium* at the terminal branches of the tree. However, we find that introgressed *S. haematobium* individuals with introgressed *S. bovis* mitochondrial genomes form two monophyletic clades. Clade “A” contains samples from Egypt, Niger, and Cote d’ Ivoire, and Clade “B” contains samples from Niger, Nigeria, and Cote d’ Ivoire; each clade spanning >1,000 Km. The most parsimonious interpretation of the phylogeny is that the introgressed *S. bovis* mitochondria share two distinct origins and imply at least two admixture events resulting from mating between a *S. bovis* female and *S. haematobium* male that occurred in the distant past. We note that laboratory crosses between *S. haematobium* are often asymmetric, and may only produce offspring when male *S. haematobium* are mated with female *S. bovis* ^40^ or produce more male offspring^31^. As females are the heterogametic sex, F1 females are expected to show reduced fitness as predicted by Haldane’s rule^41^. This may contribute to the limited number of *S. bovis* mtDNA lineages observed in *S. haematobium populations*.

Fourth, introgressed *S. bovis* nuclear loci are heavily fragmented within the *S. haematobium* genomes indicating multiple generations since introgression. Our estimates of time since introgression span 257-879 generations ago (95% confidence interval). Introgressed loci were measured in tens of kilobases (median = 76.3 Kb) and the largest blocks extended into the megabases (max = 4.05 Mb). This contrasts with early generation hybrids which would have introgressed block lengths spanning, or nearly spanning entire chromosomes^42^ which range in size from 19,481 Kb – 93,306 Kb. One Nigerian sample contained ∼25% introgressed DNA, consistent with expectations for a F2 backcross. However, the maximum introgressed fragment size in a Nigerian sample was only 2.83 Mb and median block size in these samples ranged from 47.1-97.6 Kb indicating multiple recombination events. We found that the time since introgression was significantly different between multiple countries (Figure 6). For example, neighboring countries Niger (453 generations) and Nigeria (565 generations) were not significantly different, but introgression in Cote d’ Ivoire (385 generations) appears to have occurred more recently. The variation in the estimates of generations since introgression are consistent with several regional introgression events. Alternatively, variation in age estimates between countries could reflect extrinsic factors like seasonality or intervention strategies that could lengthen or reduce generation times within sub-populations. If this were the cause, it is possible that the number of generations that have lapsed since an introgression event may vary between countries. Nigerian samples contained, significantly higher levels of introgression than other countries (Kruskal-Wallis H test statistic = 7915, *P*-value = 0.0049; Supplemental Figure 3): further analyses of Nigerian samples will be of considerable interest.

Two caveats are needed. First, our age estimates are based on the size of introgressed fragments and assume neutrality. This assumption is violated, because we see some introgressed segments are under positive selection, while others are purged resulting in introgression deserts. Violation of the neutrality assumption may add uncertainly to our age estimates. Second, given that hybridization and introgression has occurred in the past, and that hybridization can be staged in the laboratory^43^, we might expect that hybridization events may also be ongoing. One interpretation of the fragmented landscape of introgressed *S. bovis* DNA within *S. haematobium* genomes is that this results from an equilibrium between newly introgressed DNA, and selective removal (or selection for) introgressed DNA. Importantly, with both the historical and equilibrium models, the small size of introgressed fragments is clearly consistent with rare introgression, and levels of resulting interspecific gene flow are insufficient to reduce high levels of genetic differentiation between these species.

*S. bovis* shows 3.3-fold higher diversity than *S. haematobium*, while genetic diversity (π) is 2.3-fold greater in the north *S. haematobium* population than in the south African *S. haematobium* population. When the introgressed *S. bovis* alleles are removed from the analyses, this difference in genetic diversity between the north and south *S. haematobium* populations is reduced to just 1.05-fold and π is not significantly different (Figure 5B). By contrast, *F*ST values between northern and southern *S. haematobium* are consistent whether introgressed alleles are considered (*F*ST = 0.16) or not (*F*ST = 0.154) and the relationship among samples in the PCAs is nearly identical when introgressed alleles are included or excluded. These results indicate (i) that the elevated π in northern African *S. haematobium* results from *S. bovis* introgression and (ii) that northern and southern *S. haematobium* populations are well differentiated even after removing introgressed *S. bovis* alleles.

Given that introgressed *S. bovis* alleles have persisted in the northern *S. haematobium* population, we examined the data for signals of adaptive introgression. We found two introgressed genome regions with signals of positive selection in the northern population on chr four (NC_067199.1:28,348,440-28,877,530) and chr five (NC_067200.1:9,712,340-10,514,400).

Despite the convergence of signals to these regions, we were not able to identify variants driving selection in these regions. We found 37 missense SNVs where the *S. bovis* allele was nearly fixed in the northern population, but none withstood multiple test correction for directional selection. These variants occur in four genes (WormBaseParaSite v18.0^44^), a Rho GTPase-activating protein 35 (n_SNVs_ = 30; MS3_00007803) a Leishmanolysin-like peptidase (n_SNVs_ = 4; MS3_00007802), and two members of the Jumonji domain-containing protein 6 family; JMJD6_4 (n_SNVs_ = 1; MS3_00010935) and JMJD6_3 (n_SNVs_ = 2; MS3_00010934). The same Leishmanolysin-like peptidase (Table 1) has been identified as a candidate for adaptive introgression from *S. bovis* into *S. haematobium* in two previous studies^15,16^. Genes in same invadolysin gene family are known to modulate the snail host immune system in Schistosoma mansoni^45,46^ and this particular gene has been associated with cell migration and invasion in other parasitic taxa^47^.

We also observed genomic regions on three chromosomes of the *S. haematobium* samples where *S. bovis* introgression is rare or absent (Figure 2F). These introgression deserts may contain hybrid incompatibility loci that result in reduced fitness of early generation hybrids and present barriers to further introgression. Thirteen of the 19 regions occur on the scaffold representing the Z and W sex chromosomes. Eight of the deserts, including the largest occurs within the non-recombining Z-specific region of the sex chromosome. This is consistent with the purging of introgressed blocks containing deleterious alleles in non-recombining regions of the sex chromosomes^48^ which could lead to female sterility as predicted by Haldane’s rule.

Understanding hybridization and introgression between *S. haematobium* and *S. bovis* is important for disease control. If hybridization between these species is infrequent, then there may be minimal benefit in linking strategies that manage both human (*S. haematobium*) and livestock (*S. bovis*) *Schistosoma* species. Consistent with this, our results from these samples suggest that hybridization between these species is rare, and gene flow between these species is limited.

However, adaptive introgression has introduced *S. bovis* alleles into *S. haematobium* populations. This is a clear example of alleles being transferred between livestock and human parasites through introgression. Some *S. bovis* alleles have reached high frequency and are likely selectively advantageous. Future work should aim to understand how the introgressed *S. bovis* variants contribute to the fitness of *S. haematobium* individuals. The strong differentiation between northern *S. haematobium* populations, carrying introgressed *S. bovis* alleles and southern *S. haematobium* populations, with no introgression, is of particular interest. Additionally, future work should examine differences between northern and southern *S. haematobium* populations, and test whether they are reproductively isolated.

Several limitations to our study and its conclusions should be noted. First, recombination rates have not been quantified in *S. haematobium* so our estimates of age of admixture are based on recombination rates measured in *Schistosoma mansoni*^49^. To improve the accuracy of these estimates, direct measures of recombination rates from *S. haematobium* genetic crosses are needed. Second, our results pertain to *S. haematobium* and *S. bovis*. Extant hybridization between other schistosome species (*S. haematobium*/*S. guineensis* and *S. bovis*/*S. curassoni*) have been documented in field collected samples with genomic data^50,51^. Our results suggest that *Schistosoma* species pairs may form a spectrum in hybridization frequency and compatibility. Future work to understand the factors that impact hybridization and present barriers to gene flow between schistosomes species pairs will be of great interest, and can provide a more nuanced understanding of hybridization and potential implications for schistosome control.

## Online Methods

### Sample collection: description, ethics, and identification

We used samples or data from three sources. i) The first dataset was generated from samples provided by the <u>S</u>chistosomiasis <u>C</u>ollection <u>A</u>t the <u>N</u>atural History Museum^52^ which is housed at the Natural History Museum (London). SCAN samples consisted of individual miracidia and cercariae preserved on Whatman FTA cards ^53^. We analyzed 114 *S. haematobium* and *S. bovis* samples from 123 individual hosts (snails or humans) and 12 Africa countries. ii) In addition to the SCAN samples, we collected nine adult *Schistosome* worms, presumed to be *S. bovis*, from the intestines of routinely slaughtered cattle from meat vendors at three abattoirs located in Auchi, Benin City, and Enugu in Nigeria. In the laboratory, the mesenteric vessels of each purchased intestines were visually inspected for schistosome parasites. Adult schistosomes were recovered using forceps and washed in saline solution. Adult pairs were separated into males and females before being stored in 96% ethanol for subsequent DNA isolation analyses. iii) Finally, for the third source of data we used whole genome sequence data from NCBI^15,16,18,22,26,30^.

Samples provided by the SCAN repository were originally collected in accordance with protocols approved by local, state, and national authorities, including the Ministry of Health. The Imperial College Research Ethics Committee (ICREC) at Imperial College London, in conjunction with ongoing Schistosomiasis Control Initiative (SCI) activities, provided additional ethical guidance for samples collected through the CONTRAST program. Ethical clearance and study protocols for Nigerian samples were approved by the National Health Research Ethics Committee of Nigeria (NHREC) (protocol number: NHREC/01/01/2007– 30/10/2020 and approval number: NHREC/01/01/2007– 29/03/2021) and the Institutional Review Board (IRB) of University of Texas Health, San Antonio Texas, United States of America (protocol number: HSC20180612H). Informed consent was obtained from all participants, with processes tailored to ensure understanding and voluntary participation. All data were anonymized to protect participant privacy, and schistosomiasis-positive individuals were treated with a single dose of praziquantel (40 mg/kg). For livestock parasite collection, approval was secured from local veterinarians. No animals were euthanized for research purposes; *Schistosoma* samples were collected during routine activities at abattoirs. Further details on collection methods, ethical approvals, and data availability for public samples can be found in their respective publications documented in Supplemental Table 1.

Provisional species identifications were assigned to cercariae and miracidia based on sampled host. For example, miracidia hatched from eggs collected from human urine samples were assumed to be *S. haematobium* while miracidia hatched from eggs in cattle feces were assumed to be *S. bovis*. Cercariae collected from snails were identified by Sanger sequencing the mitochondrial cox1 region and the ribosomal internal transcribed spacer (ITS) rDNA region as previously described^21^. Downstream genetic analysis with whole genome SNVs was used to confirm and reassign species identifications where necessary.

### Library prep and sequencing

DNA from single parasites stored on FTA cards was subjected to whole-genome amplification (WGA) using methods previously described in ^53^. DNA was extracted from single male adult *S. bovis* worms using the DNeasy® Blood and Tissue kit before subsequent WGA. We quantified amount of schistosome DNA in each WGA sample by real time quantitative PCR (qPCR) reactions using the single copy gene α-tubulin 1 gene markers primers (*S. haematobium*: forward [GGT GGT ACT GGT TCT GGT TT], reverse [AAA GCA CAA TCC GAA TGT TCT AA]; *S. bovis*: forward [ATG GCC TCG TTA TCA ACC AT], reverse [TGG CCT CGT TAT CAA CCA TA] following previously described protocol in ^53^. DNA sequencing libraries were generated from 500 ng of DNA per sample using the KAPA Hyperplus kit protocol with the following modifications: i) enzymatic fragmentation at 37°C for 10 minutes, ii) adapter ligation at 20°C for an hour, and iii) 4 cycles of library PCR amplification. After qPCR quantification of each library with KAPA Library Quantification Kits, samples with similar concentrations were combined into pools for sequencing at 4nM, while samples with disparate concentrations were equalized in 10 mM Tris-HCl pH 8.5 before pooling. Libraries were sequenced with 150 bp paired-end reads on two Illumina NovaSeq flowcell. All resulting reads were deposited in the NCBI Short Read Archive under BioProject PRJNA636746 and are documented in Supplemental Table 1.

### Computing environment

Analyses were conducted on the Texas Biomedical Research Institute’s high-performance computing cluster, with worker nodes containing 96 cores and 1 TB of memory. Computational environments were managed using Conda v22.9.0. Environmental recipe files, Jupyter notebooks, and other code can is archived on GitHub (github.com/nealplatt/sch_hae_scan v0.1z) and at https://doi.org/10.5281/zenodo.13124719.

### Read filtering and Mapping

Raw reads were quality trimmed with trimmomatic v0.39 ^54^ using the following parameters: LEADING:10, TRAILING:10, SLIDINGWINDOW:4:15, MINLEN:36, ILLUMINACLIP:2:30:10:1:true. This command removed low quality bases at the beginning and ends of the reads, removed portions of the read where quality dropped below a minimum threshold, trimmed adapter sequences and discarded reads <36 nts. We then mapped the trimmed reads to the Egyptian-strain *S. haematobium* reference genome, GCF_000699445.3^22^, with BBMap v38.18^55^. On average the *S. haematobium* and *S. bovis* (GCA_944470425.1) genome assemblies are ∼97% similar across their genomes^18^ which should minimally affect reference biases when mapping short reads. However, to avoid reference biases we used the ‘vslow’ and ‘minid=0.8’ options with BBMap and discarded ambiguously mapping reads (‘ambig=toss’).

### Genotyping, phasing, and filtering

Mapped reads were sorted with SAMtools v1.13^56^ and checked for duplicates with GATK v4.2.0.0’s^57^ mark_duplicates. Then single nucleotide variants (SNVs) were genotyped with HaplotypeCaller and GenotypeGVCFs. To make the dataset more manageable, we genotyped each chromosome separately using the -L option. Next, we removed all indels and hard filtered SNVs based on QualByDepth (“QD < 2.0”), RMSMappingQuality (MQ < 30.0), FisherStrand (FS > 60.0), StrandOddsRation (SOR > 3.0), MappingQualityRankSumTest (MQRankSum < -12.5), and ReadPosRankSumTest (ReadPosRankSum < -8.0) with GATK’s VariantFiltration. We removed multi-allelic sites, and sites with genotype quality (GQ) <20 or read depth (DP) <8 with VCFtools v0.1.16^58^. After these filters were applied we removed genomic sites that were genotyped in ≤50% of individuals and then any individuals that were genotyped at ≤50% of sites.

SNVs on each chromosome were phased using Beagle v 5.2_21Apr21.304^59^ in windows of 20 cM and a 10 cM overlap. Currently there are no direct measures of recombination rate in *S. haematobium*. The best available data is from *S. mansoni*^49^ which has and a map length of 1134.8 cM and an estimated recombination rate (physical-to-map distance) of 244.2 Kb/cM. The genome assemblies for *S. mansoni* (GCA_000237925.5) and *S. haematobium* (GCF_000699445.3) are similar in length, 391 Mb vs 400 Mb respectively so assuming a uniform recombination rate similar to *S. mansoni* across the genome, these values are comparable to a 4.88 Mb window and a 2.4 Mb step size^49^. We used a burn in of 20 iterations and 60 iterations for the phasing run. All phased chromosome VCFs were combined into a single file using vcfcombine from vcflib v1.0.3^60^ before an additional round of post-phase filtering.

In some cases, multiple miracidia were analyzed from a single host potentially adding highly related samples to our dataset and skewing the downstream results. To remove these, we examined kinship coefficients in our samples using the autosomal chromosomes and the “– unrelated” function in king v2.2.7^61^. This parameter identifies second-degree relatives within the dataset that can be removed prior to downstream analyses. Next, we generated a set of SNVs that were common (minor allele frequency; MAF > 0.05) and unlinked. Unlinked loci were filtered with Plink v1.90b6.21^62^ by removing linked SNVs with a pairwise r^2^ > 0.2. This filter was applied in 25 Kb sliding windows with a 5kb steps. Finally, we used SnpEff v5.1^63^, to identify the impact of these SNVs on the amino acid sequence in coding regions. To do this we imported the *S. haematobium* reference genome (GCF_000699445.3) along with the associated GenBank annotations to create a custom database.

### Principal Component Analyses

We used a series of tools to explore population structure in our data sets. We used common (minor allele frequency; MAF>0.05), unlinked, autosomal SNVs and Plink v1.90b6.21^62^ to generate a principal component analysis (PCA) to examine relationships among the samples. We used a *K*-means clustering algorithm to assign each sample to between 1 and 10 populations with the Kmeans() function in sklearn.cluster v1.2.0^64^. We then used the Elbow method^65^ to examine distortion in the model and determine the optimal number of clusters in the data. Once we identified the optimal number of clusters, we assigned each sample within a cluster, and those designations were used to differentiate the *S. haematobium* populations using analyses as below. These cluster assignments were also used validate the assumed species identify of each sample.

### Admixture

We examined the ancestry of each sample with Admixture v1.3.0^25^ and the same unlinked, autosomal SNV dataset from the PCA analyses. However, we further thinned the SNV data with VCFtools v0.1.16^58^ ensuring that no two SNVs were within 10kb of each other. This step minimizes any potential effects of linkage on the results. We ran Admixture v1.3.0^25^, allowing for 2 to 20 possible population components, and used the cross-validation error to determine the optimal range^66^. Additionally, we randomly selected individuals with ≥99.999% *S. bovis* or *S. haematobium* ancestry in the k=2 analysis to serve as reference samples for each species in downstream analyses.

### Nucleotide diversity (π), Sequence divergence (*d_XY_*), and Fixation index (*F*ST)

We used scikit-allel v1.3.5 ^67^ to calculate nucleotide diversity (π), sequence divergence (*d_XY_*), and the fixation index (*F*ST) in sliding windows of 10 kb using autosomal SNVs allel.windowed_diversity(), allele.windowed_divergence, and allel.windowed_weir_cockerham_fst() functions. The weighted, Weir-Cockerham *F*ST^23,24^ was measured between species (*S. haematobium* vs. *S. bovis*) and between the *K*-means populations. Next, we used the reference panel, described above, to identify ancestry informative sites between the *S. haematobium* and *S. bovis* samples. We used scikit-allel v1.3.5’s^67^ allel.weir_cockerham_fst() to calculate *F*ST at individual sites. Only sites where *F*ST = 1 were retained.

### Biogeography

*S. haematobium* was split into two groups based on the *K*-means clustering analysis of the PCA results. At k=2 Admixture differentiated *S. haematobium* and *S. bovis* samples, but at k=3 Admixture broadly confirmed the presence of two different *S. haematobium* populations. We used the admixture proportion (Q) from k=3, to visualize how the populations were distributed across Africa. The presence of this ancestry component was extrapolated into unsampled geographic regions using the OrdinaryKriging() function implemented in pykrige v1.7.0 with a linear variogram model^68^. Geographic distances between samples were calculated with the haversine() v2.8.0 function (https://pypi.org/project/haversine/).

### Genome-wide tests for introgression

We used a series of tests to explore the presence of introgression between *S. bovis* and the *S. haematobium* populations. First, we used average_patterson_f3() from scikit-allel v1.3.5^67^ to calculate a normalized *f* ^69^ averaged across blocks of 500 SNVs. Next, we tested for gene flow using the *D*-statistic, also known as the ABBA BABA test ^70^. We used *S. margrebowiei* (GCA_944470205.2^30^) as the outgroup (O), *S. bovis* as the donor population (P3), and the *S. haematobium K*-means populations as the recipients (P1 and P2). We measured *D* across the genome in 500 SNV blocks with moving_patterson_d() in scikit-allel v1.3.5^67^. Introgressed loci were defined when *D*>0+2σ.

### Local Ancestry Assignment

For local ancestry assignment, we used RFMix v2.03-r0^28^ and TWISST v67b9a66^29^. RFMix v2.03-r0^28^ uses a random forest approach to assign local ancestry to genomic segments by comparing samples to reference panels. For this, we used the reference samples selected from the Admixture analyses. We generated a genetic map using a uniform recombination rate estimated from *S. mansoni* crosses (1 centimorgan = 287,000 bp^49^). The remainder of the parameters were set to the default.

TWISST v67b9a66^29^ uses gene trees sampled from across the genome to identify potentially introgressed loci. It does this by iteratively sampling subtrees from the gene tree and calculating relative support for each of the possible species trees. We generated gene trees from loci containing 500, phased, common (MAF >0.05) SNVs with RAxML-NG v1.1^71^. For each locus we searched for the 10 best trees and then bootstrapped the best tree for 100 replicates using the GTR+ASC_LEWIS substitution model and *S. margrebowiei* as an outgroup. Nodes supported in ≤10 bootstrap replicates were collapsed with Newick Utilities v1.6^72^. The collapsed trees were used as input for TWISST v67b9a66^29^. Samples were assigned to their *K*-means population.

### Selection

We compared selection in the *S. haematobium* intra populations using cross-population extended haplotype homozygosity (xpEHH^73^). Unphased xpEHH was measured with selscan v2.0.0^74^. The resulting unphased xpEHH values were normalized with norm v1.3.0 and the ‘--xpehh flag’. Bonferroni corrected p-values were assigned to each site. Sites with a corrected p-value < 0.01 were considered to be experiencing putative directional selection between the two *S. haematobium* populations.

### Identifying putative adaptive introgression

We searched the genome for regions with *F*ST, Patterson’s D, local ancestry, and xpEHH values indicative of adaptive introgression. To do this we examined how these values were distributed across the genome in sliding windows of 337 Kb and 3,370 bp step size; values equivalent to 1% and 0.01% of the autosomal genome. Specifically, we were looking for regions of the genome that are among the most highly differentiated between the two schistosome populations (*F*ST >= 95^th^ percentile), with statistically significant signals of introgression (Patterson’s D > 0) and directional selection (xpEHH p-value < 0.01), and the *S. bovis* alleles are at high frequency in the northern or southern *S. haematobium* populations (>95%). Patterson’s D was measured under the assumed 4-taxon tree (((southern *S*. *haematobium*, northern *S. haematobium*), *S. bovis*), *S. margrebowiei*). Windows that met these criteria were then merged together if they were within 10Kb of each other to identify loci containing signals of adaptive introgression.

### Autosomal Species Tree

To better understand the relationships among the samples, used SVDquartets^75^ as implemented in PAUP* v4.0.a.build166^76^ to generate a species tree. SVDQuartets has been shown to perform well in the presence of gene flow as was suspected here^77^. We examined 2.5m random quartets along with 100 standard bootstrap replicates. Nodes in the gene trees supported by <10% of bootstrap replicates were collapsed Newick Utilities v1.6^72^.

### Dating introgression

Recombination acts to continuously break down introgressed haplotypes. As a result, the size of introgressed haplotype blocks is directly related to the number or generations since hybridization^78^. This can be roughly estimated with the formula G=1/LP where G is generations, L is the average length of introgression haplotypes in Morgans, and P is the proportion of the genome from the major parent^79^. We identified introgressed blocks and their lengths (L) for each individual with RFMix v2.03-r0^28^ and P was estimated using Admixture (represented as *q*). A one-way ANOVA was used to identify differences in age estimates between populations (countries).

### Introgression Deserts

Some regions of the genome may be resistant to introgression. This could present as large regions lacking introgressed alleles. We used the RFMix results to identify regions of the genome where *S. bovis* ancestry was 0% in the north African *S. haematobium* populations. We log-transformed the length of each region and assigned robust Z-scores. Putative introgression deserts were regions with robust Z-scores > 3.

### Mitochondrial genome assembly and phylogeny

We used GetOrganelle v1.7.7.0^80^ to *de novo* assemble mitochondrial genomes. Specifically, we used the animal_mt model and 10 rounds of assembly with *k*-mer sizes of 21, 45, 65, 85, and 105. The mitochondrial contigs were then scaffolded with RagTag v2.1.0^81,82^ and RAxML-NG v1.1^71^ was used to generate a maximum likelihood tree of the mitochondrial genomes. We used a GTR+G substitution model to and 100 starting trees. Nodal support was assessed with 1,000 bootstrap replicates.

## Acknowledgements

We thank Sandy Smith and John Heaner for providing computational support at Texas Biomedical Research Institute’s High-Performance Computing Cluster. We would like to thank Frederic Chevalier, Winka Le Clec’h, and Kathrin Bailey for providing feedback during the course of this project. This research was funded by the National Institute of Allergy and Infectious Diseases (NIAD R01 AI166049-01). Schistosomiasis Collections at the Natural History Museum (SCAN) was supported by Wellcome Trust grant 1045958/Z/13/Z. The field work referenced in this study relied on various organizations and funding bodies across multiple countries. In Angola, funding was provided by the Calouste Gulbenkian Foundation in collaboration with the Centro de Investigação em Saúde de Angol, the Programa Nacional de Doenças Tropicais Negligenciadas and the Ministério da Saúde de Angola. In Côte d’Ivoire and Niger, the research was funded by SCORE through the Bill & Melinda Gates Foundation, RISEAL-Niger, Université Félix Houphouët-Boigny, and the Centre Suisse de Recherches Scientifiques en Côte d’Ivoire. The Eswatini National Control Program, Environmental Health Unit, and the School Health Program contributed significantly to the work in Eswatini. The CONTRAST project in Kenya, Uganda, and Zambia was funded by the European Commission, the Kenya Medical Research Institute, National Museums of Kenya, Uganda Ministry of Health, and the University of Zambia. In Liberia, the UK Department for International Development provided funding via SCI (now Unlimit Health) and the Ministry of Health of Liberia. Madagascar’s efforts were funded through SCAN, Unlimit Health, and the Madagascar Ministry of Public Health. Work in Namibia was funded by the END Fund and the Namibia Ministry of Health and Social Services. In Zanzibar, the ZEST project was funded by SCORE, the Ministry of Health (Zanzibar), and the Public Health Laboratory—Ivo de Carneri, Pemba.

Individual collectors are listed in the Supplemental Table 1.

## Author Contributions

Roy N Platt II: Conceptualization, Formal Analysis, Writing – Original Draft, Writing - Review & Editing. Elisha Enabulele: Conceptualization, Formal Analysis, Investigation, Resources, Writing – Original Draft, Writing - Review & Editing. Ehizogie Adeyemi: Resources. Marian O Agbugui: Resources. Oluwaremilekun G Ajakaye: Resources. Ebube C Amaechi: Resources. Chika E Ejikeugwu: Resources. Christopher Igbeneghu: Resources. Victor S Njom: Resources. Precious Dlamini: Resources. Grace A. Arya: Investigation. Robbie Diaz: Investigation. Muriel Rabone: Resources, Writing - Review & Editing. Fiona Allan: Resources, Writing - Review & Editing. Bonnie Webster: Resources, Writing - Review & Editing. Aidan Emery: Resources, Writing - Review & Editing. David Rollinson: Resources, Writing - Review & Editing. Timothy J.C. Anderson: Conceptualization, Formal Analysis, Writing – Original Draft, Writing - Review & Editing, Supervision.

## Corresponding authors

Correspondence to Roy N. Platt II or Timothy J. C. Anderson.

---

**Supplemental Figure 1.**
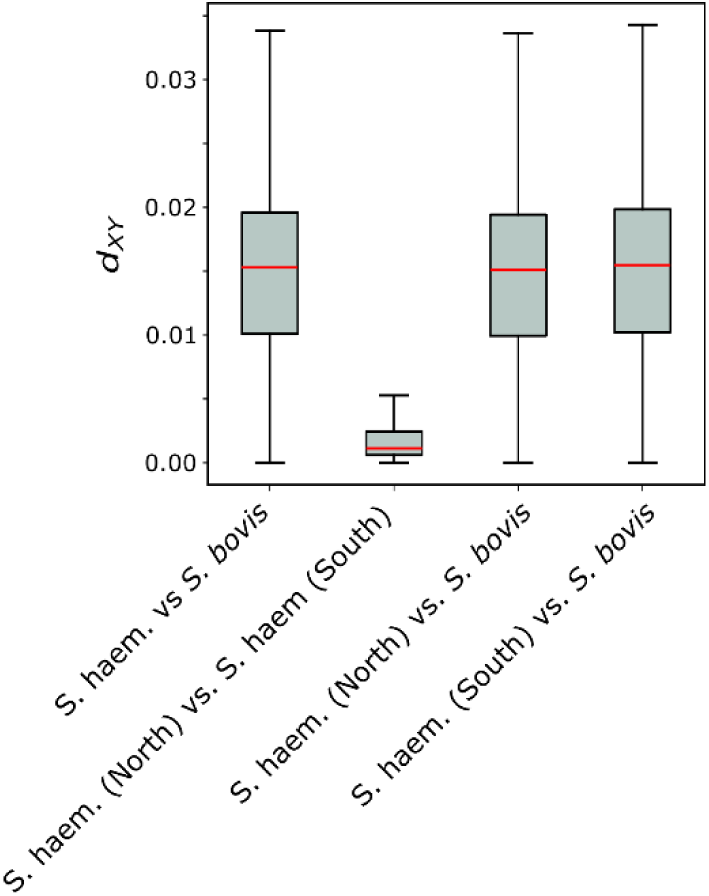
Sequence divergence (*dXY*) between *S. bovis* and *S. haematobium*. We examined *dXY* across the genome in 10Kb windows. Sequence divergence was the same across all comparisons between *S. haematobium* samples and *S. bovis* regardless of the population tested (*dXY* means = 0.0144-0.0148). By comparison sequence divergence between northern and southern *S. haematobium* populations was nearly 7x lower (mean *dXY* =0.002).

**Supplemental Figure 2.**
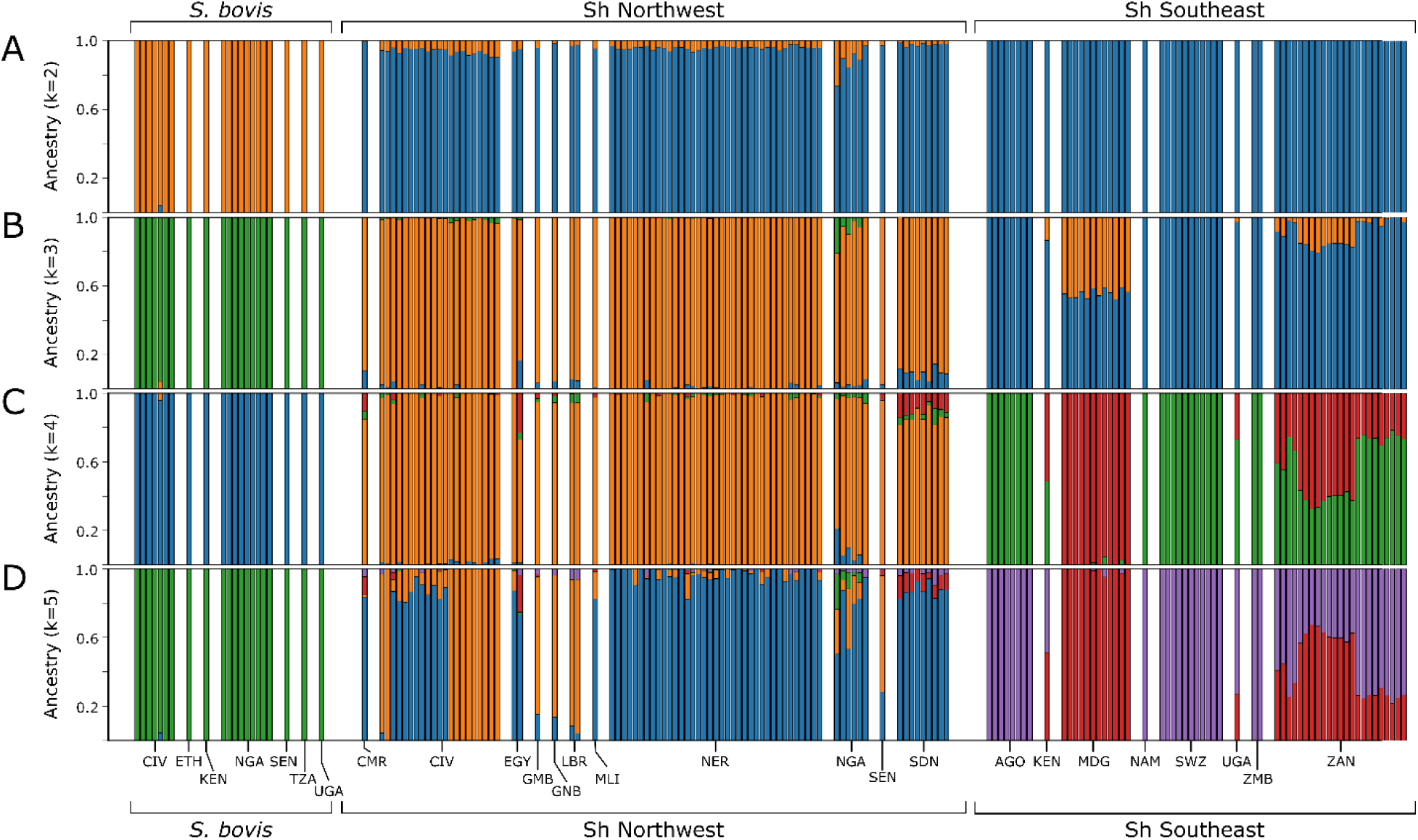
Whole genome ancestry assignment with Admixture. We examined multiple different population (k) sizes with Admixture. (A) At k=2, *S. haematobium* and *S. bovis* were separated, and two general populations were identified within the *S. haematobium* samples corresponding to a northern and southern population. We also partitioned the data into (B) three populations (k=3) and (C) four populations. (D) Five populations (k=5) was the optimum number according (Evanno *et al*. 2005). Here the samples show clear distinctions between the two *S. haematobium* populations. Country Codes are as follows: “AGO”: Angola, “CMR”: Cameroon, “CIV”: Cote d’ Ivoire, “EGY”: Egypt, “SWZ”: Eswanti, “ETH”: Ethiopia, “GMB”: Gambia, “GNB”: Guinea Bissau, “KEN”: Kenya, “LBR”: Liberia, “MDG”: Madagascar, “MLI”: Mali, “NAM”: Namibia, “NER”: Niger, “NGA”: Nigeria, “SEN”: Senegal, “SDN”: Sudan, “TZA”: Tanzania, “UGA”: Uganda, “ZMB”: Zambia, “ZAN”: Zanzibar.

**Supplemental Figure 3.**
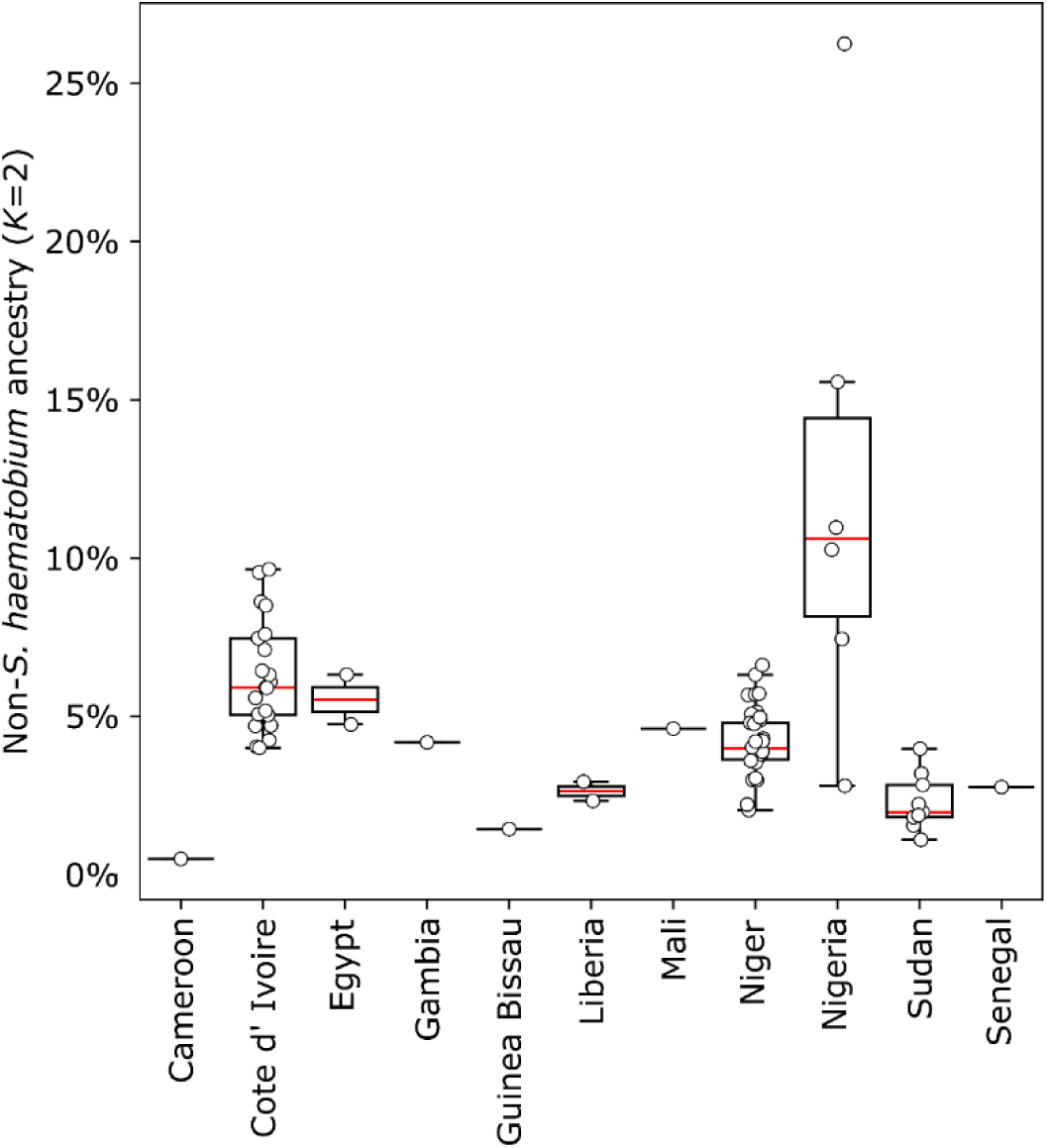
Comparison of non-*S. haematobium* ancestry calculated from Admixture (k=2) in each north African sample. – Ancestry of each sample was assigned to up to two different population components with Admixture. These two components were maximized in samples from southern Africa or S. bovis samples. By comparison, north African samples were a composite of these two populaitions with low, but varying levels fo the *S. bovis* compponent found in each individual. The population component corresponding to *S. bovis* ancestry was significanly higher in Nigerian that in other north African countries (Kruskal-Wallis H test statistic = 7.915, P-value = 0.0049).

**Supplemental Figure 4.**
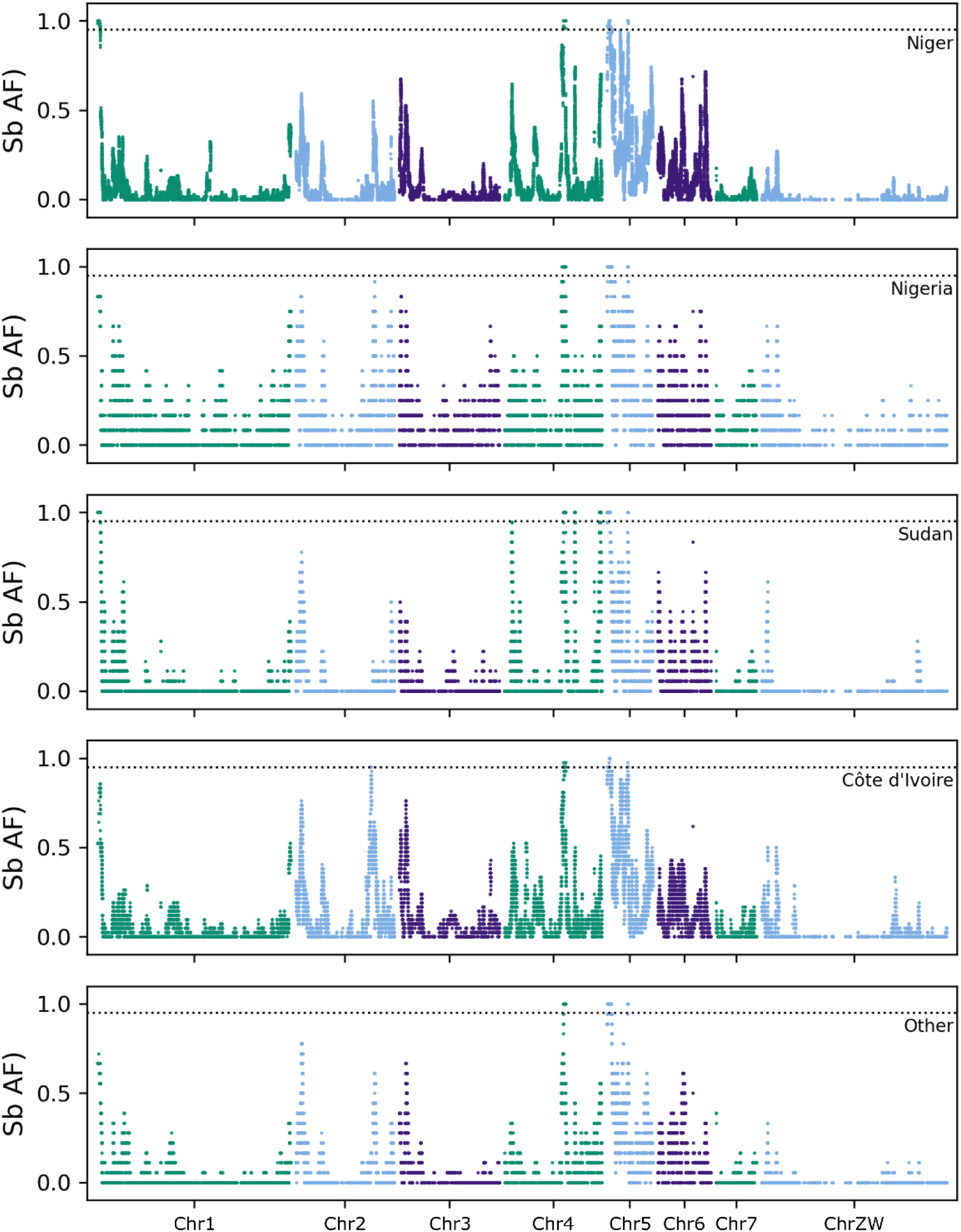
*S. bovis* allele frequency across the genome within *S. haematobium* samples from Northern African countries. The frequency of *S. bovis* ancestry across the genome is shown for each of the northwest African countries. In general, the distribution of *S. bovis* alleles is similar for each population. This consistency is an indicator of historic introgression event(s). The dotted line indicates 95% allele frequency.

**Supplemental Figure 5.**
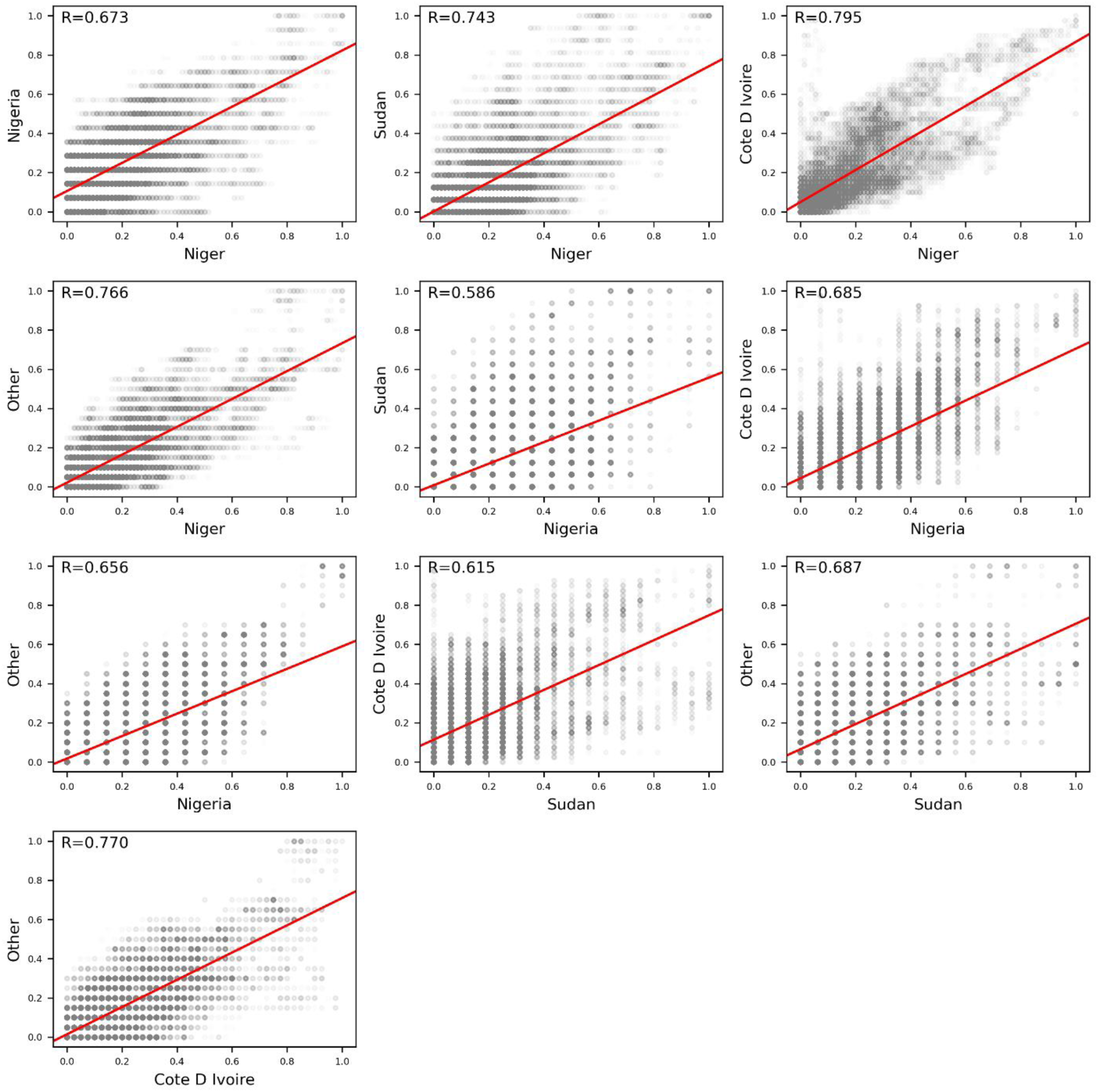
Pairwise comparison of introgressed *S. bovis* allele frequencies within northern *S. haematobium* samples by country. Introgressed *S. bovis allele* frequency is positivley correlated between countries. Pearson’s correlation coefficient (*R*) is >0.586 in all comparisons. The correlation of introgressed allele freuqencies between populations up to 3,338 Km apart is consistent with older introgression events. .

**Supplemental Figure 6.**
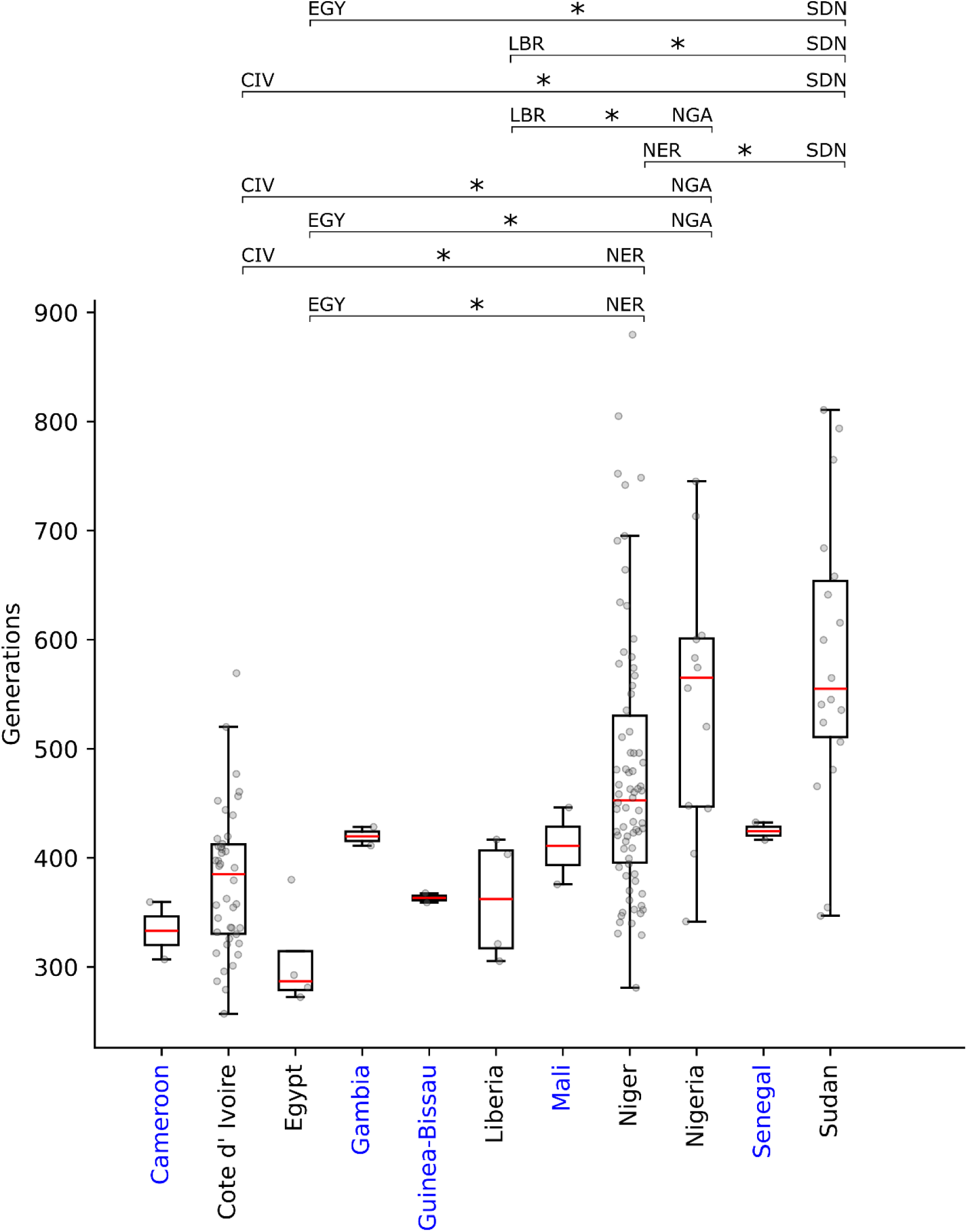
Estimated number of generations since admixture with *S. bovis*. We estimated the number of generations since admixture for each sample in the northern *S. haematobium* population by examining the length of introgressed *S. bovis* loci with in the genomes. Individual estimates for each sample for each country are shown as two grey points, one for each haplotype. Results from a one-way ANOVA indicated that age estimates varied significantly between countries. Countries with a single individual (two haplotypes) are shown in blue and were not included in the ANOVA analyses. A “*” indicates p-values < 0.05. Differences in ages may indicate multiple introgression events.

